# Hypertrophic cardiomyopathy disease results from disparate impairments of cardiac myosin function and auto-inhibition

**DOI:** 10.1101/324830

**Authors:** Julien Robert-Paganin, Daniel Auguin, Anne Houdusse

## Abstract

Hypertrophic cardiomyopathies (HCM) result from distinct single-point mutations in sarcomeric proteins that lead to muscle hypercontractility. While different models account for a pathological increase in the power output, clear understanding of the molecular basis of dysfunction in HCM is the mandatory next step to improve current treatments. An optimized quasi-atomic model of the sequestered state of cardiac myosin coupled to X-ray crystallography and *in silico* analysis of the mechanical compliance of the lever arm allowed us to perform a systematic study for a large set of HCM mutations. We define different classes of mutations depending on their effects on lever arm compliance, sequestered state stability and motor functions. This work reconciles previous models and explains how distinct HCM mutations can have disparate effects on the motor mechano-chemical parameters and yet lead to the same disease. The framework presented here can guide future investigations aiming at finding HCM treatments.

---

Hypertrophic cardiomyopathies (HCM) are the most prevalent inherited cardiac diseases, affecting one individual per 500^1^. Clinical manifestations range from asymptomatic to mechanical or electrical defects leading, in the most severe cases, to heart failure or sudden death^2^. Virtually all HCM is caused by mutation of genes necessary for cardiac muscle contraction and at least 80% of familial HCM is caused by mutation of two sarcomeric proteins: MYH7 (ventricular β-cardiac myosin) and MyBP-C3 (cardiac myosin binding protein C)^3^. More than 300 mutations in the β-cardiac myosin heavy chain gene are known to cause HCM in adults and mutations causing severe childhood HCM have also been described^1,4^. Although HCM mutations are found in all regions of β-cardiac myosin, clusters of mutations in regions such as the converter are known to be particularly severe^5^. The motor cycle of myosins couples ATP hydrolysis to actin-based force production upon the swing of its distal region called lever arm, including the converter. The motor undergoes critical conformational changes and several structural states of different affinity for F-actin and different lever arm positions have been described^6^ (Fig.1A). To date, the effects of HCM mutations on motor activity have not been precisely predicted^7^. While recent advances give hope in future treatments of such a disease with small molecule modulators of myosin function^8,9,10,11,12^, it is essential to progress in understanding of the impact of β-cardiac myosin mutations to guide future therapeutic strategies.

**Figure 1:**
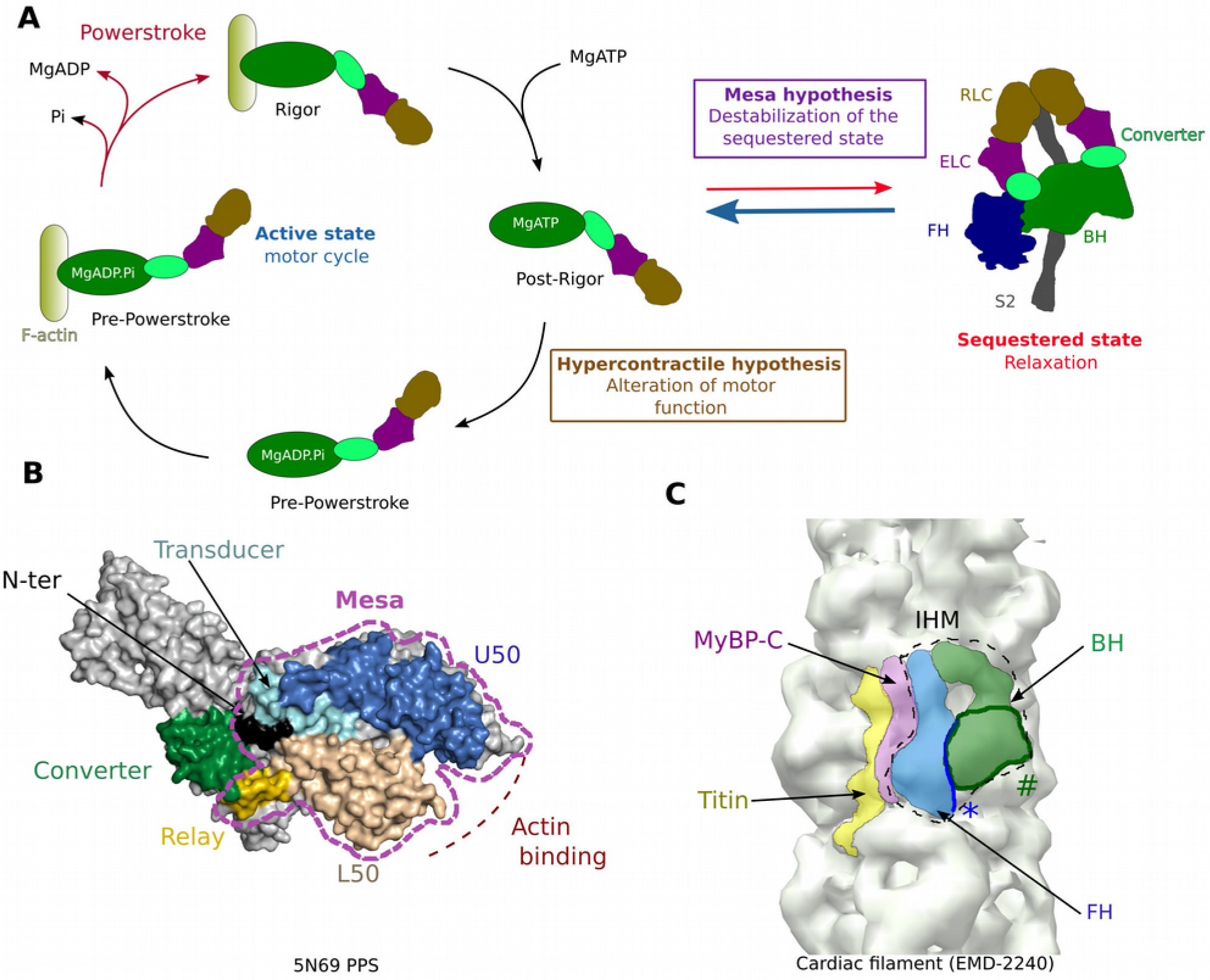
Myosin mesa and sequestered state. (A) Schematic representation of the motor cycle and the regulation of the β-cardiac myosin activity. On the left, when the motor detaches from the track upon ATP binding, the motor adopts the post-rigor (PR) state in which the lever arm is down and the motor has poor affinity for F-actin. During the recovery stroke, repriming of the lever arm leads to the pre-powerstroke (PPS) state in which hydrolysis can occur. The swing of the lever arm (powerstroke) upon reattachment of the motor to F-actin is coupled with the release of hydrolysis products. The nucletotide-free or rigor state has the highest affinity for F-actin. On the right, scheme of the sequestered state that is formed during relaxation. According to the mesa hypothesis, HCM mutations disrupt the sequestered state, increasing the number of myosin heads available to produce force. According to the Brenner hypothesis, HCM mutations alter the motor activity of the myosins. (B) The mesa (purple dashed lines) is a long and flat surface of the myosin head composed of several myosin subdomains and that is conserved in myosin IIs. (C) Electron density map of the human cardiac filament obtained by negative staining^2,3^ (EMDB code EMD-2240). In the relaxed state, interactions between the blocked head (BH) and the free head (FH) of the myosin 2 dimer stabilize an asymmetric configuration. Extra densities of the filament correspond to other components of the thick filament, the cardiac myosin MyBP-C and the titin. Location of the mesa for each head indicates that the FH mesa (*) interacts with the BH while the BH mesa (#) is buried and interacts with components of the thick filament.

Brenner and co-workers have been the first to propose a mechanism of HCM development^13,14,15^ based on HCM mutations located in the converter, a motor subdomain critical for the swing of the myosin lever arm during force production^16^. They assessed that these mutations alter the mechano-chemical parameters of individual myosin heads thus modifying the power output of the cardiac muscle^13,14,15^ (Fig. 1A). According to Brenner’s results, HCM converter mutations alter the stiffness of the lever arm and thus modify the intrinsic force produced by each head, highlighting the essential role of the converter for motor compliance^14^. However, how a single point mutation in the converter can alter the global compliance of the molecular motor has yet to be described.

Recently, a contrasting hypothesis has been proposed by Spudich and co-workers who found that HCM mutations are strongly enriched on a flat surface of the myosin head, the ‘mesa’^17,18,19^ (Fig. 1B). The mesa hypothesis^17,19^ proposes that mutations of the mesa and converter regions trigger HCM by increasing the number of active heads since they lead to the destabilization of the myosin sequestered state, in which the heads are inactive and unable to interact with actin (Fig. 1A). In cardiac filaments, the sequestered state would be a way to regulate the number of active heads available to produce force during contraction^19^. In this state, the two heads of a myosin molecule form an asymmetric dimer (Fig. 1A) first described for the off state of smooth muscle myosin^20^, which was later identified in myosins of invertebrate and vertebrate striated muscle, including ventricular cardiac muscle^21,22,23,24^. The widespread occurrence of the ‘interacting heads motif’ (IHM) suggest a conserved mechanism to sequester these different myosins (Fig. 1C). In vertebrate muscles, such as cardiac, the sequestered state has been proposed to be the structural basis of a “super-relaxed” functional state that occurs during relaxation with extremely low ATPase activity^25,26,27,28^. The transition between the sequestered and the active states is driven by several parameters such as calcium concentration, interaction with MyBP-C and mechanical load^29^. Current low resolution structural models indicate that several HCM mutations could weaken the so-called IHM^30,19^ thereby increasing the number of active heads. Recent data is consistent with this conclusion since the myosin intrinsic force is little affected by adult-onset HCM converter mutations^31^ (Fig. 1A). Current models of the IHM lack sufficient resolution to describe the interfaces of the IHM at an atomic level^19^ and how mutations can weaken or abolish the sequestered state. A higher resolution structural model is thus required. In the context of HCM, the main challenge is to identify how different mutations can impair the cardiac myosin activity or its regulation and lead to the HCM disease.

Here, the cardiac myosin converter/essential light chain interface is described for the first time from a 2.33 Å crystal structure of β-cardiac myosin. Novel insights on the role of the converter in myosin compliance and in stabilization of the sequestered state are provided from *in silico* studies of four severe HCM mutations located in the converter. In order to analyze how HCM mutations may destabilize the IHM or affect motor activity, an optimized molecular model of the sequestered state was built using our high resolution cardiac myosin structures. Our results provide precise insights for the development of the HCM pathology, explaining how HCM mutations can modify motor activity and compliance of single heads and how they may also increase the number of heads available for force production by destabilizing the sequestered state.

## Results

### High resolution Structure of the β-cardiac myosin head - description of the converter/ELC interface

We determined the 2.33 Å resolution structure of the Bovine Cardiac Myosin S1 fragment (BCMS1) complexed with MgADP in the post-rigor (PR) state (PR-S1), a myosin conformation populated upon ATP-induced detachment of the motor from actin (Fig. 2A, Supp. Table 1). The same fragment was previously crystallized in the PPS state complexed with *omecamtiv mecarbil*^11^ (OM-PPS-S1, PDB code: 5N69) (Supp. Fig. 1B). The ELC bound to the IQ motif was built *ab initio* from well-defined electron density (Supp. Fig.1A).

**Figure 2:**
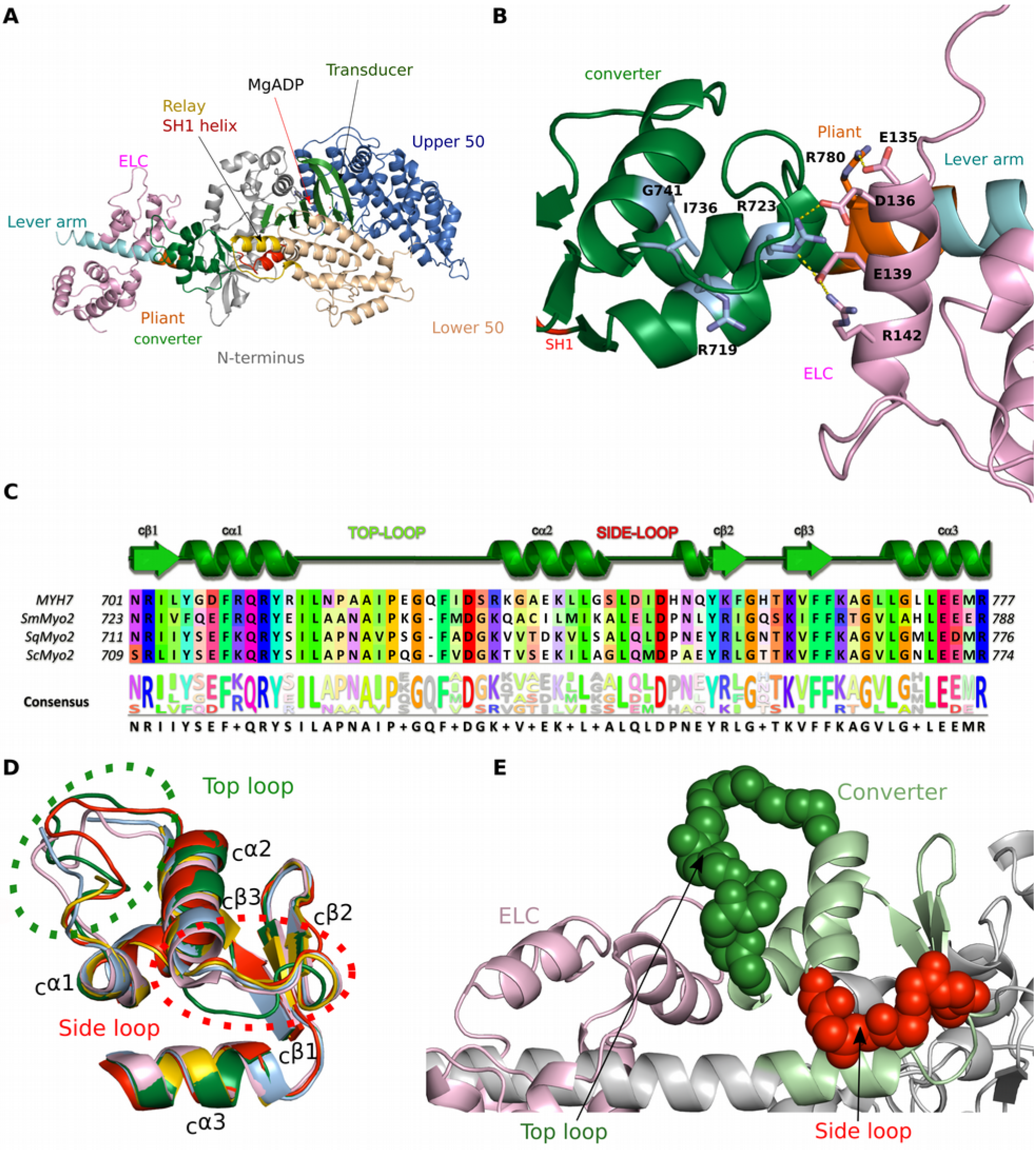
Crystal structures of β-cardiac myosin and description of the converter/ELC interface. (A) X-ray structure of β-cardiac myosin S1 complexed with MgADP in the post-rigor (PR-S1) state. **(B)** Interface between the converter and the ELC as found in the PR-state. This interface involves mainly electrostatic interactions between a cluster of negatively charged residues of the ELC (D136; E135 and E139) and positively charged residues of the heavy chain (R723 and R780). Side chains of interacting residues (sticks) and polar interactions (yellow lines) are represented. Four converter mutations studied in this work (R719W; R723G; I736T; G471R) are colored in light blue. **(C)** Structure alignment of converters of the Myosin 2 superfamily. MYH7: bovine (*Bos taurus*) β-cardiac myosin; ScMyo2: bay scallop (*Argopecten irradians*) Myosin 2; SqMyo2: Longfin inshore squid (*Dorytheutis pealeii*) Myosin 2; SmMyo2: chicken (*Gallus gallus*) gizzard smooth muscle myosin 2. **(D)** Superimposition of several Myo2 converter structures: from PR-S1 colored in green; from SqMyo2 (PDB code 3I5F, pink); from SmMyo2 (PDB code 1BR1, red); from ScMyo2 in the PPS (PDB code 1QVI, light blue) and Rigor (PDB code 1SR6, yellow) states. **(E)** Location of the top loop (green) and the side loop (red) in the cartoon representation of the crystal structure of the β-cardiac myosin PR-S1 (grey) with the converter colored in light green and the ELC colored in light pink. The top loop is part of the converter/ELC interface.

These structures thus allow the description, for the first time, of the critical ELC/converter interface for two different positions of the cardiac myosin lever arm. There is no major variation in the lever arm coordinates of the PR and the PPS structures (rmsd 0.98 Å) (Supp. Fig. 1C). The pliant region (R777-783) at the junction between the converter and the IQ motif/ELC^16^ is straight in both structures (Supp. Fig. 1C). In the two structures, interactions between the ELC and the converter are mainly electrostatic (Fig. 2B, Supp. Fig. 1B) via a network of charged residues that contribute to the stabilization of the lever arm.

The overall fold of the converter is similar to that found for smooth and invertebrate striated muscle myosins, with the exception of two loops displaying sequence variations, that we define as the ‘*top loop*’ and the ‘*side loop*’ (Fig. 2C, 2D, 2E). The side loop appears to assist stabilization of the fold of the converter, while the top loop forms a part of the converter/ELC interface. Thus, these structures provide complete atomic coordinates that are essential to evaluate the compliance of the lever arm and to study the consequences of specific missense mutations in this region.

### The cardiac lever arm internal dynamics

The dynamics within the cardiac myosin lever arm were studied in 30 ns molecular dynamics experiments (see methods) using coordinates from the PR-S1 lever arm structure corresponding to the converter (701-777) and 1^st^ IQ motif (778-806) as well as the bound ELC. We computed the r.m.s. fluctuation during the simulation for all Cα atoms and represented it with the “putty representation”^32^(Fig. 3). The WT simulation reveals strong internal dynamics of the converter (rmsd of Cα between 0.6-5.8 Å) linked to large fluctuations in the top and side loops (Fig. 3A). The converter/ELC interface does not behave as a stable supramodule. In fact, dynamic fluctuations promote formation of labile polar bonds depending on the relative orientation of the ELC and converter. Some residues such as E732 from the converter (_C_E732) and R783 from the pliant region (_P_R783) increase the dynamics at the converter/ELC interface by oscillating between partners and thus favoring rearrangement. For example, the top loop residue CE732 oscillates between CR723 and ELCR142 by a mechanism analogous to musical chairs: a single negative charge (CE732) alternates between two nearby positive charges (CR723 and ELCR142) (Fig. 3A). These labile bonds regulate the conformation and dynamics of the top loop yielding controlled dynamics at the converter/ELC interface. In addition, the space explored by the top loop is also controlled by the side chain of CR719 interacting with carbonyls of the top loop in certain conformations that are labile due to the top loop dynamics. In contrast, the stable interaction of PR780 with the ELC (ELCD136), constrains the dynamics of the pliant region^16^, controlling the myosin lever arm deformability. We studied *in silico* R719G and R780E mutations to test the importance of such anchoring residues. These mutations result in drastic increase in the amplitude of the converter dynamics via destabilization of contacts at the pliant region (Supp. Fig. 2A, 2B, 2C). Large lever arm compliance resulting from such mutations would likely affect motor activity. This study thus highlights that the dynamics in this multi-domain lever arm is controlled by crucial bonds within the lever arm. However, the lever arm is dynamic rather than rigid and modulation of the conformations of the top loop of the converter as well as of the pliant region greatly contribute to the dynamics at the converter/ELC interface.

**Figure 3:**
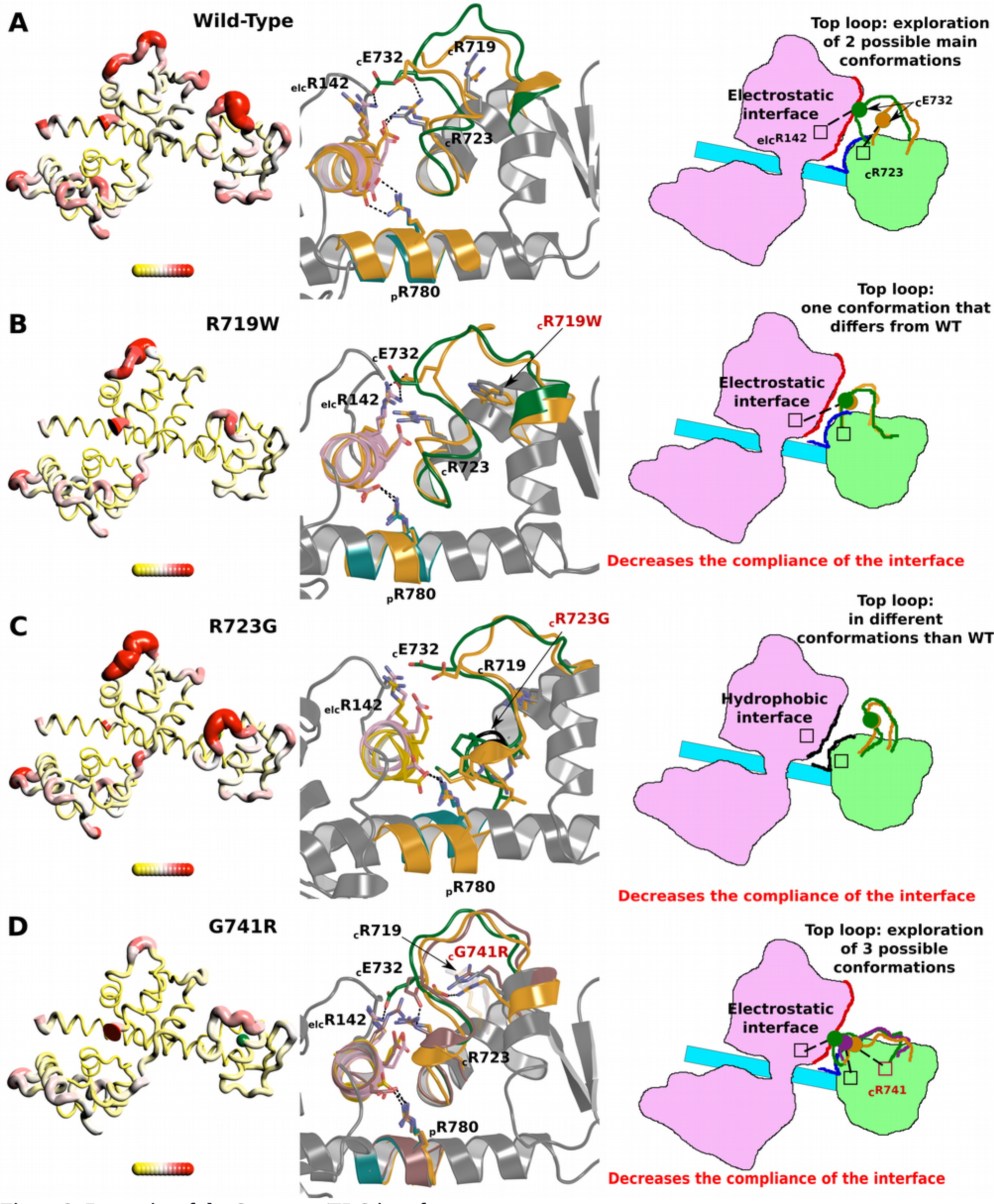
Dynamics of the Converter/ELC interface: Schematic representation of the results from the molecular dynamics simulations for the wild-type (WT) and the three HCM mutants that have been analyzed *in silico*: R719W, R723G, G741R. On the left, a “Putty representation” of the β-cardiac myosin converter and first IQ (aa 701-806) bound to the ELC. R.M.S. fluctuations during 30 ns simulations are represented with r.m.s. scale ranging from 0.6 Å (in yellow) to 5.8 Å (in red). In each structure, the position of the mutated residue is represented by a green sphere. On the center, the structure of the interface between the ELC, the converter and the pliant region is represented as well as the position of key-residues maintaining the interface and its plasticity. On the right, a schematic representation of this region is displayed. The different populations of the top loop allowed by the dynamics of this region are drawn and the nature of the interactions between the converter and the ELC is indicated. **(A)** WT, **(B)** R719W, **(C)** R723G, **(D)** G741R.

### Converter HCM Mutations alter the lever arm internal dynamics via long-range effects on a global charge network at the converter/ELC interface

Differences in the lever arm dynamics linked to HCM mutations may result in effects in the intrinsic force produced, the sensitivity of the motor to external load as well as the formation of the sequestered state (see below). We next searched with simulations how the controlled dynamics observed in the WT lever arm are affected by severe HCM converter mutations. We focused on three converter mutations for which single molecule studies had been performed. The intrinsic force was virtually unchanged by the G741R mutation and was only modestly (15-30%) decreased for R719W and R723G^31^ (Supp. Fig. 3). An additional effect of these mutants on myosin regulation cannot be excluded since some HCM mutations have already been reported to disrupt interfaces of the IHM^33^. *In silico* studies of these mutants show a large effect on the top loop conformation which results in reduction of the lever arm pliancy.

The R719 side chain interacts transiently with the top loop and helps in maintaining its conformation (Fig. 3A). The **R719W** mutation has two effects, first there is a destabilization of the top loop conformation found in WT and second, the contacts promoted by the bulky W719 decrease the internal dynamics of the converter and of the converter/ELC interface (Fig. 3B). The R723 side chain is part of the network of labile converter/ELC interactions that facilitate dynamics in the WT lever arm (Fig. 3A). The **R723G** mutation removes a charged residue crucial for this interface as well as for the stabilization of the top loop conformation. Interestingly, in this mutant, the ELC/converter interface is defined via formation of hydrophobic interactions and the top loop is not part of this interface. While the internal dynamics of the R723G converter is locally high due to destabilization of the top loop, drastic loss of the controlled “musical chairs” dynamics at the converter/ELC interface leads to more stiffness overall (Fig. 3C). The **G741R** mutant introduces a charged and bulky side chain in the second helix of the converter which results in an extra charged partner at the converter/ELC interface that disrupts the musical chairs dynamics found in WT (Fig. 3D). Thus, the top loop _C_E732 residue can now interact with this mutant R741 side chain and is no longer confined to oscillations between _C_R723 and _ELC_R142. This leads to a shift in the position of the top loop that reduces the internal dynamics of the G741R converter and the ELC/converter interface compared to WT (Fig. 3D). Overall, these simulations show that although the lever arms of these HCM mutant myosins are slightly stiffer or similar to WT in terms of compliance, the mutations have a large effect on the top loop conformation and its dynamics and result in disruption of the dynamic network at the converter/ELC interface. The consequences on force production are likely moderate since the lever arm remains able to transmit and amplify the internal conformational changes of the motor. This is consistent with what has been measured previously, since the effect of these mutations on fibers mechano-chemical parameters remain modest^31^.

We also studied a fourth HCM mutant^34^ that affects a top loop residue, I736T^34^ (Supp. Fig. 2D). Only local changes in the top loop conformation result from this mutation leading to limited consequences on the dynamics of the converter or the interface with the ELC. To reveal how I736T can result in HCM, it is thus critical to describe the effect of the I736T mutation on the sequestered state. We thus further our analysis by describing how HCM mutations affect the formation of the β-cardiac myosin sequestered state.

### Sequestered state model and the role of the converter/ELC interface

In the sequestered state, myosin heads are in a conformation with the lever arm primed, which is a hallmark of the PPS state^20,24^. Therefore, the cardiac PPS structures we have determined provide the best starting model to date to generate a high resolution *in silico* model of the cardiac myosin sequestered state (see Methods). By a combination of homology modeling, ambiguous docking methodology^35^, molecular dynamics and fit in the two existing maps^23,30^ (see Methods), we have built a model of the β-cardiac myosin sequestered state that can provide a much improved description of the inter-head interactions responsible of its stabilization (Fig. 4A). The model perfectly fits in current low-resolution available maps (Supp. Fig. 4). This model is the best to date for two reasons (see Methods): **(i)** it is based on a high resolution PPS state structure^11^ (PDB code 5N69) that is precise enough to ensure the correct position of motor domain residues; **(ii)** the use of molecular dynamics allowed us to model realistic inter-head interactions and to ensure that our final model displays good geometrical parameters. Thus using the latest results in electron microscopy, we provide a quasi-atomic model of the sequestered state that is a powerful tool to study how mutations may affect the stability of this sequestered state.

**Figure 4:**
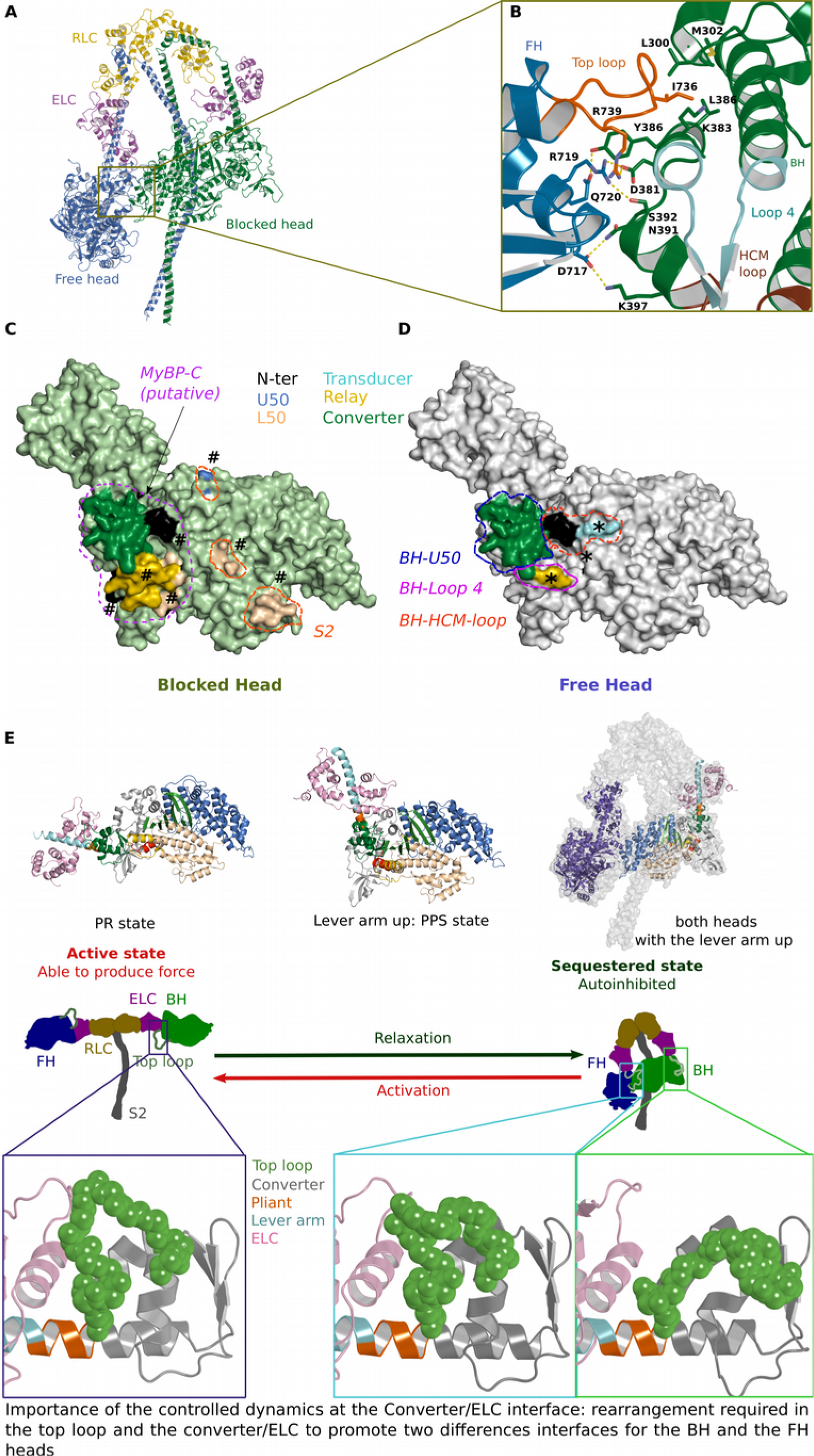
An optimized model of the sequestered state of β-cardiac myosin. **(A)** β-cardiac myosin sequestered state modeled from the cardiac myosin S1 PPS structure. This model results from optimization of the intra-head interactions that occur upon formation of the IHM (Woodhead et al., 2005) with two asymmetric heads: the free head (FH) and the blocked head (BH). **(B)** Detailed analysis of a region of the interface involving the FH-converter and the BH-U50 subdomain. The FH-converter is a major component of this interface, in particular the top loop plays a major role. **(C)** Surface of the FH-mesa and the FH-converter involved in interactions with the BH. **(D)** Surface of the BH-mesa and the BH-converter involved in interactions with the FH. A putative surface of interaction with the partner MyBP-C is also represented. The regions of the interface that are part of the mesa are labelled with a ‘*’ and a ‘#’ on the FH and on the BH respectively. **(E)** Schematic representation analyzing the prerequisites to form the IHM. On top, different myosin conformations illustrate the differences in position of the lever arm in the PR and the PPS states. Two heads in the PPS conformation are shown in the sequestered state. On the center, a schematic representation of the two states adopted by the heavy meromyosin (HMM) is shown. On the bottom, the different conformations of the top loop present in the PR crystal structure and in the BH and the FH of the model are displayed in cartoon representation. To form the sequestered state, the lever arm has to adjust their conformation differently in the two heads, in particular for the top loop. Note however that the pliant region remains close to the conformation of the WT PPS structure. Dynamics at the ELC/converter interface of the two heads of the myosin dimer is thus important to promote sequestration of the heads.

In the new quasi-atomic IHM model, consistent with previous models^30,33^ (PDB code 5TBY; MS01), the sequestered state is asymmetrical and composed of two folded heads (Fig. 4A). The S2 coiled-coil interacts with the mesa of the so-called blocked head (BH) (Fig. 4A, 4C). The BH is unable to interact with actin since a part of its actin binding interface is docked on the free head (FH) via the flat surface mainly composed of the FH-mesa and FH-converter (Fig. 4B, 4D). We mapped on the surface of the mesa and the converter of the FH and the BH the regions that are involved with inter-or intra-molecular interactions (Fig. 4C, 4D). The FH head interacts with the BH head, while the BH-converter putatively interacts with MyBP-C. The BH/MyBP-C interface is supported by previous experiments^33^ and the architecture of the cardiac filament^23^. There are two main differences in the sequestered model we propose compared to previous models. **(i)** The motor domain coordinates are much improved and reliable since the molecular dynamics have optimized the overall geometry of the model and the intra-molecular interactions that occur upon head/head recognition; **(ii)** The lever arm differs drastically in the pliant region for both heads. In previous models, the use of the smooth myosin structure (1BR1) as a template had imposed a bend at the pliant region (Supp. Fig. 5A), which is not observed in the cardiac myosin PPS X-ray structure.

This model provides atomic-level description of inter-head interactions in the sequestered state. A remarkable feature of the FH/BH interface is the major role of the FH-converter and specifically the top loop: (i) I736 (FH top loop) occupies a hydrophobic pocket of the BH-U50 in a lock-and-key fashion (Fig. 4B, Supp. Fig. 5E), (ii) polar and charged residues from the FH converter, including R719 (converter) and R739 (top loop), create an extended network with the BH-U50 (Fig. 4B). These interactions significantly differ from the previous models in which the top loop is in a drastically different conformation and was not predicted to form contacts that would mobilize I736 (Supp. Fig. 5E).

Finally, this model highlights a role in the lever arm dynamics for the formation of the sequestered state (Fig. 4E). In relaxing conditions, formation of the interactions between heads requires that both heads adopt an asymmetric conformation, in particular in the hinges of their lever arm, while the two motor domains position the lever arm up (PPS state). Interestingly, the converter/ELC interface and in particular the conformation of the top loop is different in the BH and the FH heads. This implies that controlled dynamics at the converter/ELC interface is important for the establishment of the inter-head interfaces and thus mutations affecting the dynamics can impair the formation and/or the stability of the sequestered myosin state.

### Implication of the cardiac IHM model in disease

The atomic model of the sequestered state built from high resolution cardiac structures and optimization of the intra-molecular contacts provides a powerful structural framework to investigate the impact of HCM mutations. We have carefully studied a set of 178 mutations previously described as triggering the HCM phenotype (www.expasy.ch, ^18,30^; see Supp. Table 2). We evaluated the effects of each of these mutations based on the structural myosin models that define how myosin can produce force^6^ and how it is sequestered for inactivation (Table 1, Supp. Table 2). This analysis resulted in a classification of the mutations in six classes that provide an overview of the different molecular consequences of HCM-causing mutations that can trigger the disease (Fig. 5A, 5B). Each proportion evoked in this section is calculated as a percentage of the entire set of 178 selected mutations.

**Figure 5:**
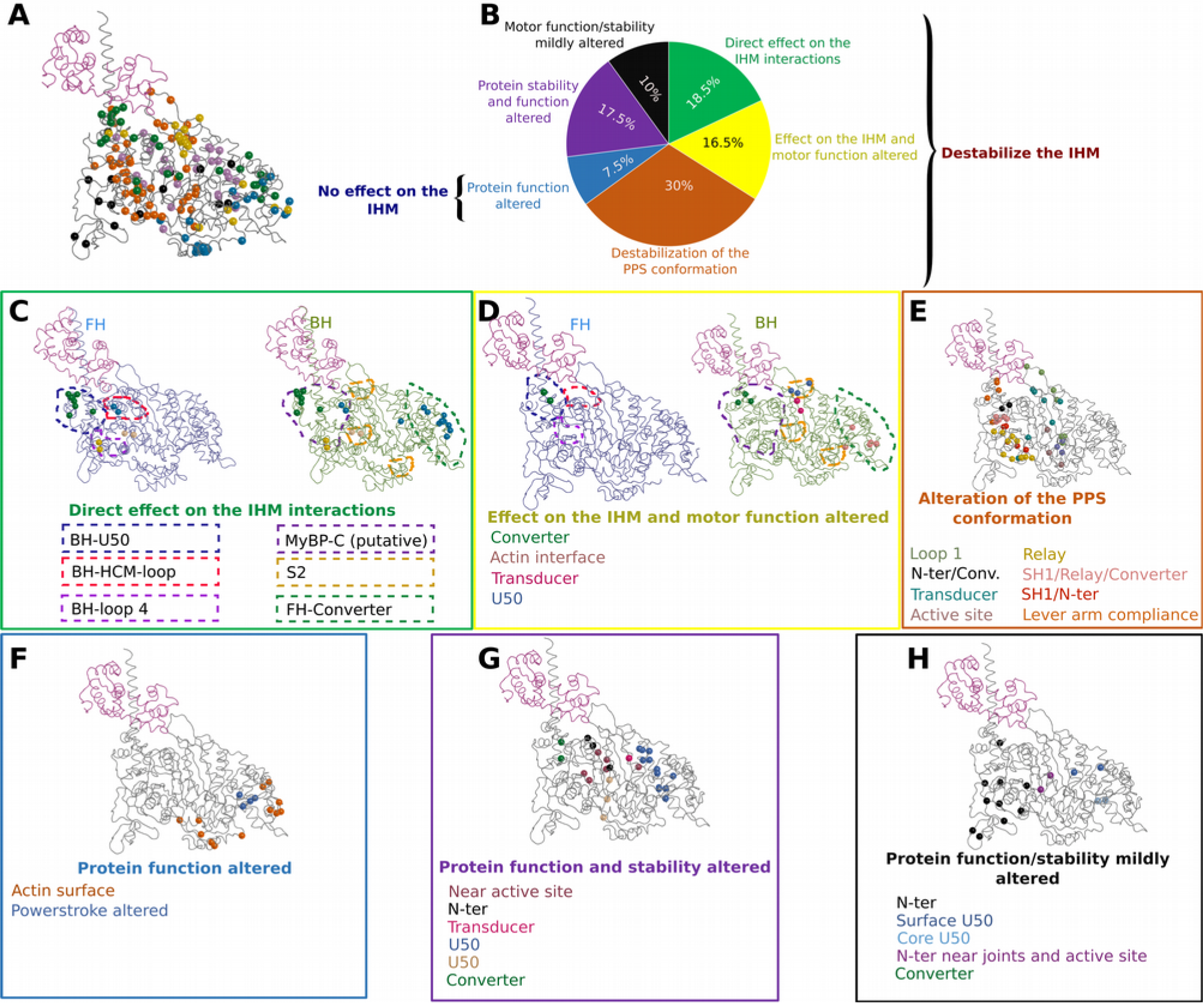
structural and functional consequences of HCM mutations. **(A)** represent the HCM mutations positions (balls on the crystal structure of the PPS state in ribbon). **(B)** A chart pie represents the proportion of mutations belonging to six classes depending on their effect on the structure, the function and the stability of the IHM. These six classes are: mutations destabilizing the IHM (green); mutations destabilizing the IHM and the motor function (yellow); mutations destabilizing the PPS conformation and the IHM (orange); mutations that alter the protein function (blue); mutations that alter the protein function and the protein stability (purple) and mutations that mildly affect protein function or stability (black). **(C), (D), (E), (F), (G)** and **(H)** show the distribution of mutations on the myosin motor domain structure for each class. The balls are colored depending on the effect and the position of the mutations. For classes affecting directly the IHM (**C**, **D**), the surfaces of interaction are represented in dotted lines.

**Table 1.**
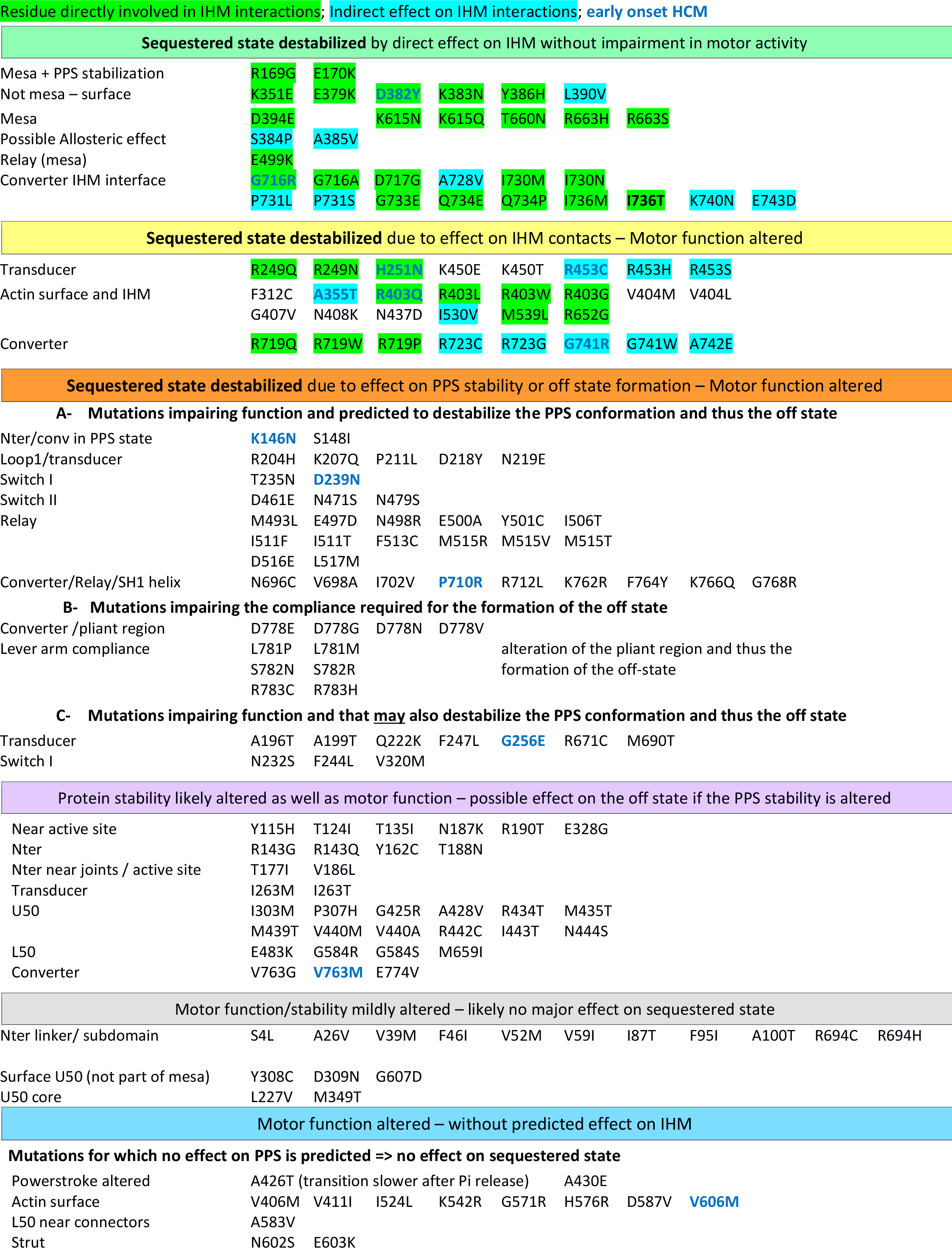
Classification of a set of 178 HCM mutations depending on their effects.

As previously proposed, some mutations are predicted to affect the IHM stability since they impact residues found at the interfaces that stabilize the off state^33,30^ (Table 1, Fig. 5C, 5D, 5E). Among these mutations, 18.5% of the set represent mutations that destabilize the IHM with no significant effect on the motor function (Fig. 5B, 5C). Some mutations that directly affect the IHM stability also have effect on motor function (Fig. 5B, 5D, 16.5% of the set). These mutations are located in the transducer, the actin binding interface or the converter which are regions of interest in the mechano-chemical cycle of the motor (Supp. Table 1). Interestingly, a large number of mutations affecting the sequestered state (30% of the set) are not directly located at the interfaces of the IHM but are predicted to alter the stability of the PPS conformation that is necessary to form the IHM (Fig. 5E, Table 1). In this case, the effects of the mutations are diverse, they can affect regions important for allosteric communication during the motor cycle (Switch I, Switch II, relay or SH1 helix), destabilize interactions between regions necessary to maintain the PPS state or alter the compliance of the lever arm at the pliant region (Fig. 4E) required for the IHM formation.

In addition, this analysis defines three other classes of mutations that impact the motor function and/or its structural stability (35% of the set). Among these, an important class regroups mutations with no effect on the IHM stability while strong effects on motor function are predicted (Fig. 5B, 5F). This class that regroups 7.5% of the set is particularly interesting since it predicts that alteration in motor function rather than on the number of heads available for contraction can also lead to HCM. One of the mutations, A426T, occurs at a position that has been described in Myosin VI (A422L) to slow the transition after Pi release^36^. This mutation indeed introduces a bulky residue in the 50 kDa cleft that must close during the powerstroke to bind actin strongly. This indicates that some of these mutations can lead to HCM by altering purely the states and transitions populated during the powerstroke. Another set of mutations is predicted to alter significantly both motor function and its stability (Table 1, Fig. 5G, 17.5% of the set). Since mutations from this class are predicted to affect the stability of the motor, we cannot exclude that some of these mutations may result in decrease stability of the PPS and consequently on the IHM. Finally, we defined a last class (Table 1, Fig. 5H) with 10% of the mutations for which only mild alteration of the local structure or the function of the protein is predicted. These mutations are mainly located in the N-terminus and their functional study under load is required to better delineate their effect on motor function.

Altogether, our results show that HCM mutations can have a wide variety of effects. Indeed, these mutations are found in all the subdomains of the myosin heavy chain (Fig. 5A, Supp. Table 2) and interestingly close mutations in space can belong to distinct classes and induce very different effects. This illustrates the complexity of the HCM pathology and the fact that HCM mutations can have not only effects on the IHM but also on motor function. While awaiting further biophysical measurements (currently limited to a small set of mutations^31,37,38^ (Supp. Table 2)), the classification of these mutations using structural models will open new avenues to investigate how altered motor function and/or its regulation can lead to Hypertrophic Cardiomyopathy.

## Discussion

### The role of the converter/ELC interface for the lever arm compliance

Molecular dynamics allowed us to describe the key-role of the top loop and of electrostatic interactions in maintaining and controlling the converter/ELC interface dynamics. Here we highlight that loss of this controlled dynamics can occur from multiple mutations at the converter/ELC interface or at the pliant region, which corresponds to the highly deformable hinge point between the converter and the IQ/ELC modules of the lever arm^16^. In this regard, dynamics in the lever arm integrate from all interactions that either regulate the converter fold stability or the interactions that form between the converter and IQ/ELC modules of the lever arm.

The *in silico* study of a set of converter mutations highlights the previously unanticipated controlled dynamics that naturally occur between the converter and the ELC which must be sufficiently rigid for myosin activity but also sufficiently pliant to allow the formation of the sequestered state of β-cardiac myosin. Thus this study not only supports the previous assumption that the converter is essential in maintaining the proper stiffness of the lever arm^13^; it indicates how the inherent complex interplay between residues of the Converter/ELC can be easily dysregulated. Interestingly, a different point mutation at a particular position (R719W and R719G) can have antagonistic effects on the lever arm pliancy. This illustrates how the sequence of this region finely tunes the lever arm dynamics and validates the *in silico* approach to investigate the effect of mutations on the dynamics of this region. The converter HCM mutants that we analyzed have two effects (i) they modify the conformation of the top loop and (ii) they slightly increase (R719W, R723G, G741R) or have no effect (I736T) on the stiffness of the lever arm. This agrees with previous single-molecule measurements that determined that R719W, R723G and G741R have no major effect on force production of S1 fragments^31^ (less than 30% decrease) (Supp. Fig. 3).

### Deciphering the molecular consequences of HCM mutations

The optimized quasi-atomic sequestered model (Fig. 4A) we built from atomic structures of the cardiac head allows us to predict the consequences of a set of 178 HCM mutations and demonstrates that this disease may occur from either dysregulation of the motor activity itself or the dysregulation of the sequestered state that shuts down myosin activity. Our results support the previous hypothesis that some of the HCM mutations can disrupt the IHM and thus increase the number of heads available to participate in force production^17^. This is the case for at least 65% of the set we examined (Fig. 5B). While some HCM mutations can directly affect the contacts within the IHM, other mutations are predicted to affect the IHM stability (Table 1), in particular by altering the stability of the PPS state of the myosin heads, which is the structural state adopted by both heads in the sequestered state (Fig. 4E). Several mutations can have dual effects; it is the case for example of the extensively studied R403Q located in the so-called HCM-loop involved in actin binding. Single-molecule studies report that the R403Q mutant has a decreased affinity for actin in the strongly-bound states and displays mechanical defects^38^. While the motor function is affected^38^, this mutation likely also weakens the IHM since the HCM loop of the BH interacts with the N-ter subdomain and transducer of the FH (Fig. 4D). Our analysis indicates that for several mutations, such as R403Q, the outcome will be an increase in heads participating in contraction with an altered mechanical function. In addition, our results clearly explain the consequences of the selected set of HCM converter mutations studied in molecular dynamics. These mutations destabilize the sequestered state by three mechanisms (i) they all affect the dynamics and the conformation of the top loop which is a part of the inter-head interface; (ii) at least two of them (I736T and R719G) change a residue which is directly involved in the interface and (iii) three of these mutations (R719W, R723G and G741R) decrease the dynamics of the converter/ELC interface which is necessary to form the sequestered state since the conformation of this interface is different in the FH and in the BH compared to the active state (Fig. 4E).

An unanticipated result from this study is the description of an interesting class of mutations (7.5% of the set) that effects motor activity without any predicted effect on the IHM stability. Interestingly, previous work proposed that HCM mutants can be differentiated from other cardiac pathologies since they would mainly disrupt the IHM while Dilated Cardiomyopathies (DCM) mutations would mainly alter myosin function, often being located close to the active site^30^. According to our results, this statement has to be nuanced since more than 7.5% of the HCM mutations we have studied are also predicted to only alter myosin function. In addition, some HCM mutations in this class involve positions close to the active site (for example Y115H and T124L). DCM mutations are predicted to have a strong effect in diminishing the power output of cardiac contraction (Sup. Table 3). Future studies on cardiac myosin under load^12^ are essential. In particular such studies will distinguish whether mutations located in the N-terminus (Fig. 5H) may alter the activity of this motor under load. Recent studies have indeed highlighted a role for the N-ter subdomain of myosin-V and myosin-Ib in the transitions associated with the powerstroke and the force sensing mechanism^39^. Important future functional studies are required to define more precisely the frontier between HCM and DCM at a molecular scale. In particular, it will be critical to test whether the HCM mutations located close to the active site, in the 50 kDa cleft or at the actin binding interface (Fig. 5F) result in gain of power output of cardiac contraction, in contrast to DCM mutations which are predicted to result from a deficit in motor function^30^ (Sup. Table 3). Studies of atypical mutations that lead to different heart impairment will also be of particular interest to investigate how the fate of cardiac cells depend on specific loss of myosin function. This is the case for the R243H mutation which leads to an apical HCM and DCM disease, or of three DCM mutations which are predicted to directly destabilize the IHM (Sup. Table 3).

### Implications for future treatments

Recently, the discovery of small molecules able to modulate the activity of β-cardiac myosin have provided new pharmaceutical perspectives for treatment of cardiac diseases^8,9,10,11^. The mechanism of action of a cardiac myosin activator, *Omecamtiv mecarbil* (OM) has been elucidated: OM favors the PPS conformation and thus increases the number of heads able to participate in force production upon the systole on-set^11^. Interestingly, a recent study has concluded that OM is not compatible with the sequestered state of cardiac myosin while the myosin inhibitor blebbistatin (BS) favors the sequestered state^40^. Our results are consistent with this study since the site of blebbistatin is internal and is not affected by IHM contacts. Thus, BS binding is compatible with the sequestered state. The fact that BS favors the off state^40^ is a strong argument in favor of the model previously proposed^41^ from low resolution EM maps which we have defined at a better resolution here with both heads mainly unaffected for the motor domain but which uses hinges at the converter/ELC interface to allow the asymmetric association of the myosin heads to form the IHM (Fig. 6). In contrast, the binding site of OM is located in a pocket between the converter and the N-terminal subdomain^11^ which restricts the compliance of the converter required for the formation of the sequestered state (Fig. 6). Thus, stabilization of the pre-powerstroke state of the motor is not sufficient to favor the myosin sequestered state and shutting down of motor activity during relaxation. Discovery of drugs that directly affect or favor the formation of the IHM motif would be of exquisite use to control more precisely the motors available.

**Figure 6:**
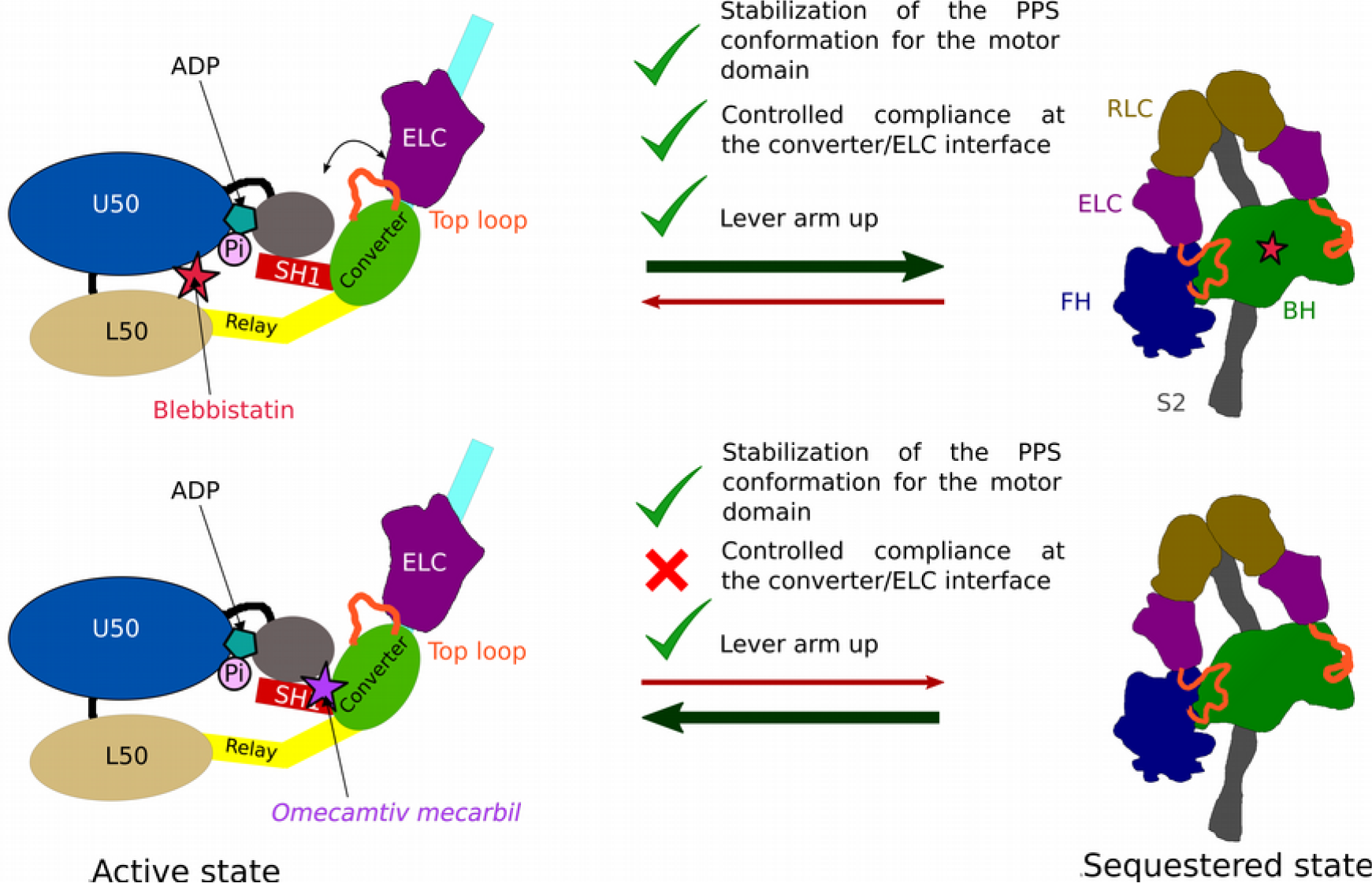
antagonistic effects of *omecamtiv mecarbil* (OM) and blebbistatin (BS) on the sequestered state. Schematic representation of the effects of the inhibitor BS and the activator OM. On top: BS occupies a pocket within the motor domain core, close to the active site. Occupation of this pocket by BS allows to stabilize the PPS conformation but it has no effect on the local compliance of the lever arm required to form the sequestered state. Thus BS binding is compatible with the IHM and favors the sequestered state. On the bottom: OM occupies a pocket at the interface between the converter and the N-ter subdomain^11^. This pocket is expected to decrease the compliance of the converter/ELC interface and constrain the lever arm in a particularly primed position. There is thus a loss in the compliance required to form the sequestered state if OM is bound to the heads of the Myo2 dimer. This explains why OM is incompatible with the formation of the sequestered state.

In this work, we indicate that HCM mutations can have different effects on the motor activity and on the regulation of cardiac myosin. It is thus unlikely that a unique drug can be used to treat all classes of HCM. Critical functional studies are required to identify how different mutations lead to impairment and how drugs modifying myosin activity or the stability of the PPS state may correct the effect of genetic mutations that belong to different classes. Since some HCM mutations may also impair the binding site of myosin drug modulators, it is critical to investigate how diverse specific drugs can restore myosin activity upon binding to diverse pockets of the myosin motor. The current knowledge about the rearrangements within the myosin motor upon force production^6^, the development of precise methodologies to study these mutations^42,31,12^ and the first *in vivo* studies about how drug may slow the progression of HCM disease^10^ announce that research will bring new therapies for this disease in the near future.

## Material and Methods

### Protein purification

Bovine cardiac fragment S1 has been purified from fresh heart as described in ^11^.

### Crystallization, data collection and processing

Bovine cardiac fragment S1 (25 mg/ml) crystallized at 4°C by the hanging drop vapor diffusion method from a 1:1 mixture of protein, 2 mM MgADP and precipitant containing 12.5% PEG mix MMW; 10 % glycerol; 7.5 % Sodium-Tacsimate pH 6.0; 3.3 % DMSO; 0.5 mM TCEP. PEG mix MMW was prepared by mixing stock solutions (50%) of PEG 2K, 3350, 4K and 5K MME in equal volume^43^. Crystals were transferred in the mother liquor containing 25% glycerol before flash freezing in liquid nitrogen. X-ray diffraction data were collected on the ID23.2 beamline at the ESRF synchrotron (Grenoble). Diffraction data were processed using the XDS package^44^. The crystals belong to the P1 space group with two molecules per asymmetric unit. The cell parameters and data collection statistics are reported in Supp. Table 1.

### Structure determination and refinement

Molecular replacement was performed with human cardiac myosin (residues 3-706) without water and ligand (PDB 4P7H) with Phaser^45^ from the CCP4i program suite. Manual model building was achieved using Coot^46^. Refinement was performed with Buster^47^. Structure determination indicated that this crystal form corresponds to a post-rigor state. The final model has been deposited on the PDB (PDB code 6FSA)

### Model building and isoforms

In this work, we chose to use the bovine isoform of the β-cardiac myosin. This choice has been done because all the S1 fragment crystal structures to date are from bovine isoform (PPS, PDB code 5N69 and PR, PDB code 6FSA). Both isoforms are very close and share 98% identity on the heavy chain (MYH7) and the differences are not located at the interfaces of the IHM.

### Sequestered state model

Two molecules of OM-PPS-S1 (PDB code and one molecule of cardiac S2 (2FXO) have been fitted in the 20 Å resolution electron density map obtained from Cryo-EM of thick filament of tarantula (*Aphonopelma sp*.; EMDB code EMD-1950) *Flexor Metatarsus Longus* striated muscle^48^. Fitting in electron density was performed with the function “fit in map” of USCF Chimera^49^. The RLC (bovine MYL2) and the second IQ motif were homology modeled based on squid isoforms (PDB code 3I5F) and fitted in the electron density. The model was made continuous and prepared with the CHARMM-GUI^50^ with the Quick MD Simulator module and the CHARMM36^51^ force field. The model was improved with several iterative steps of docking between the two heads performed with HADDOCK ^35,52^ and molecular dynamics without constraints (30 ns) on the complete model with Gromacs^53^. The only stage where interacting constraints were used was the HADDOCK docking and alternate possible assemblage modes were considered at this time. This docking stage thus allowed us to identify the best IHM model which was compatible with the low resolution EM map.

The same procedure has been followed to fit our model in the 28 Å resolution electron density map from EM data of human cardiac thick filament^23^ (EMDB code EMD-2240).

In previous studies, overall features of the sequestered state have been described from EM structures of tarantula leg striated muscle at a resolution of 20 Å, the best resolution to date^24,48^ (PDB code 3JBH) and from those of human cardiac muscle at 28 Å^23^ (EMDB code EMD-2240). Even if the thick filaments from these muscles have different helical geometries, the EM map and the structure of the two interacting heads are similar, although the resolution is not sufficient to discuss precisely the interfaces between the two heads. The sequestered model from tarantula striated muscle (PDB code 3JBH) has been obtained by a combination of homology modeling with the smooth muscle myosin sequestered state as a template (PDB code: 1l84), with flexible fitting and minimization in order to reduce steric clashes^24^. Recent models of the cardiac sequestered state have been obtained by homology modeling from the tarantula model, minimization and rigid body fit in tarantula and cardiac maps^33,30^ (MS01 available on spudlab.stanford.edu; PDB code 5TBY). However, these models are not precise enough to discuss specifically the impact of a point mutation in an interface.

The molecular model we have built differs from those previously proposed MS01^33^ and 5TBY^30^. The main difference is that our model has been computed from crystal structures describing the motor domain head at atomic resolution (PPS bound to *Omecamtiv mecarbil*, PDB code: 5N69) and that molecular dynamics was used to refine the interfaces between the two heads. We are consequently confident that the modeled interactions between the FH and the BH correspond to a realistic and optimized interface for the IHM. This refined model is the only one to date to highlight the role of the top loop (in the FH head) for its interaction with the BH head. No bend in the lever arm is present for this refined model unlike what was previously proposed in other models (based on the previous use of the PPS atomic model of smooth muscle myosin (1BR1) in which a sharp bend occurs at the pliant region (Supp. Fig. 5C) possibly due to a crystal packing artefact). To favor IHM intra-molecular interactions, a bend was introduced at the junction between the first and the second IQ (Supp. Fig. 5A). The lack of atomic resolution structures for the cardiac RLC bound to the second IQ makes this region of the model likely the least accurate. It has been modelled starting from a homology model of SqMyo2 (PDB code 3I5G). In the final steps of modeling, the S2 fragment was added. While the structure of this region is known, no precise information exists for its orientation: this is thus also a part of the model that would benefit in the future from higher resolution data. In summary, the sequestered state model was built from atomic resolution crystal structures of the motor domain that allowed us to describe refined molecular contacts between the two myosin motor domains after fitting of the model in the available EM maps and optimization of interactions (Supp. Fig. 4A, 4B).

### Molecular dynamics for the lever arm simulations

Molecular dynamics simulation inputs were prepared with the CHARMM-GUI^50,54^ with the Quick MD Simulator module. The CHARMM36 force field^51^ was used to describe the full systems. Gromacs^53^ (VERSION 5.0-rc1) was used to execute the simulations of 30 ns each. Each molecular dynamics simulation has been carried out multiple times (at least twice) in order to be sure that the results are reproducible.

### HCM and DCM mutations picking and selecting

We analyzed a set of 178 HCM mutations (Table 1, Supp. Table 2) and 24 DCM mutations (Supp. Table 3) occurring in the motor domain of β-cardiac myosin heavy chain (MYH7). Mutations were initially picked in expasy database (www.expasy.ch) and in recent publications^18,30,33^. For each mutation, we carefully checked the original publication that described the mutation in order to be sure that the mutation was strictly related to an HCM or a DCM diagnosis. Percentages are calculated relative to the entire set of 178 HCM mutations that we analyzed.

## Acknowledgements

We thank James J. Hartman for providing the protein (Bovine cardiac fragment S1) and beamline scientists of id23.2 (ESRF) for excellent support during data collection. We also thank Margaret A. Titus, Karl J. Petersen, Olena Pylypenko and H. Lee Sweeney for critical reading of the manuscript. J.R.-P. was the recipient of an Association Française Contre les Myopathies (AFM) fellowship 18423. A.H. was supported by grants from CNRS, FRM DBI20141231319, AFM 17235. The A.H. team is part of the Labex CelTisPhyBio:11-LBX-0038, which is part of the IDEX PSL (ANR-10-IDEX-0001-02 PSL).

## Contributions

J.R.-P., D.A. and A.H. designed the research; J.R.-P. crystallized and solved the crystal structure; D.A. performed the modeling and molecular dynamics; A.H. classified the mutations and analyzed the consequences; all authors discussed and analyzed the data; J.R.-P., D.A. and A.H. wrote the manuscript.

---

**Supplementary Figure 1:**
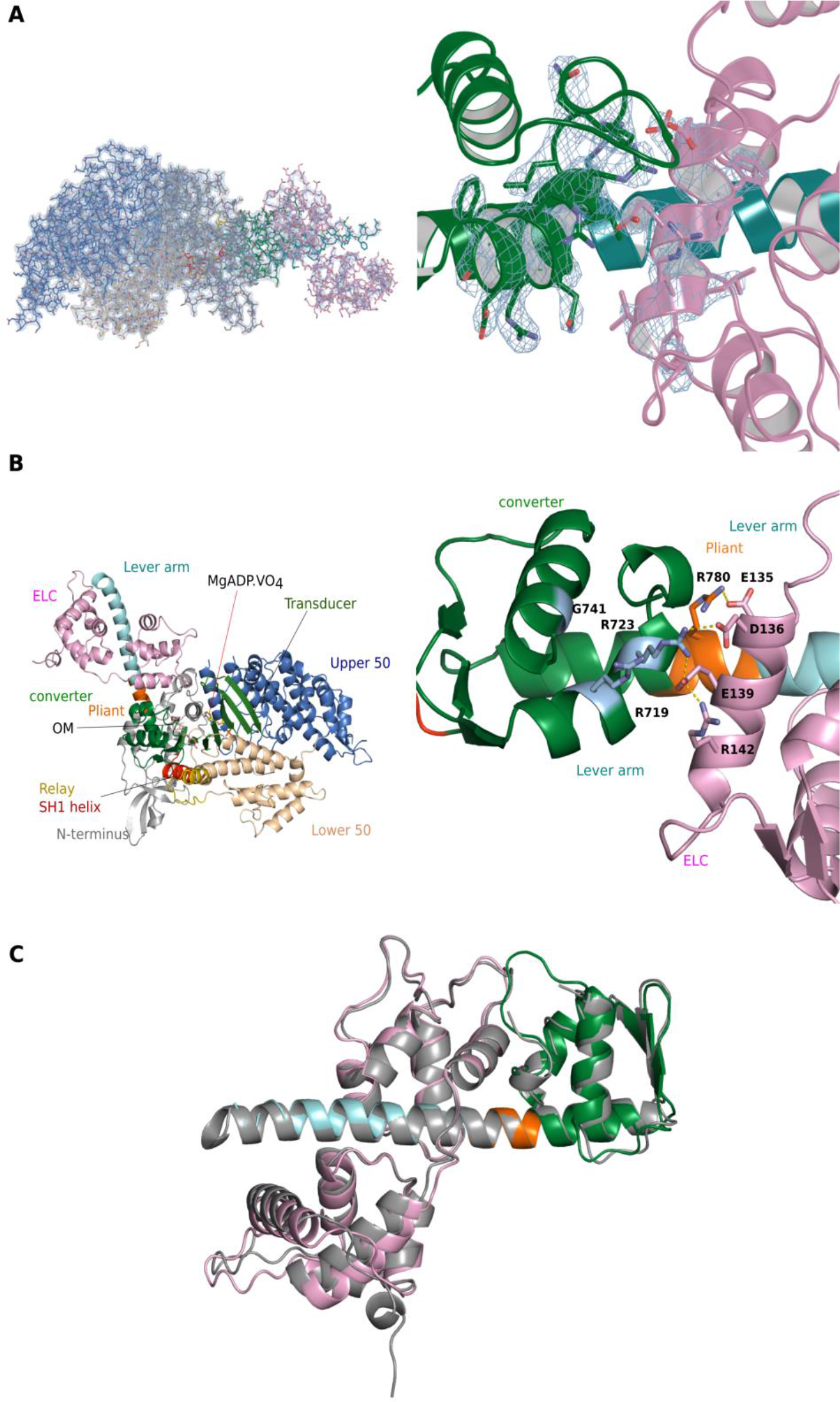
Electron density map of the PR-S1 crystal structure and the ELC/converter interface. **(A)** On the left, electron density map of the crystal structure of the β-cardiac myosin S1 fragment in the post-rigor state (PR-S1). The electron density map in pale cyan corresponds to a 2Fo-Fc map with a contour of 1.0 σ. The different subdomains of the model (stick representation) are colored: N-terminus (grey); U50 (marine blue); L50 (wheat); relay (yellow); SH1 helix (red); converter (green); lever arm (cyan); ELC (light pink). On the right, electron density map (2Fo-Fc; contour of 1.0 σ) for the converter/ELC interface. Residues involved in the interface are well defined in electron density and the side chains can be clearly identified. The ELC backbone is clearly defined as well as the side chains of the E helix that interacts with the converter. **(B)** On the left, X-ray structure of β-cardiac myosin S1 complexed to the activator *omecamtiv mecarbil* in the pre-powerstroke state (PDB code: 5N69). On the right, interface between the converter and the ELC as found in the PPS-state. **(C)** Cartoon representation of the lever arm region (converter, pliant, IQ motif and ELC) of β-cardiac myosin in the PR state (same color code as in Fig. 2A and 2B) superimposed with that found in the PPS state structure (in grey) (PDB code: 5N69). Both structures superimpose on this region with a r.m.s.d. of 0.982 Å.

**Supplementary Figure 2:**
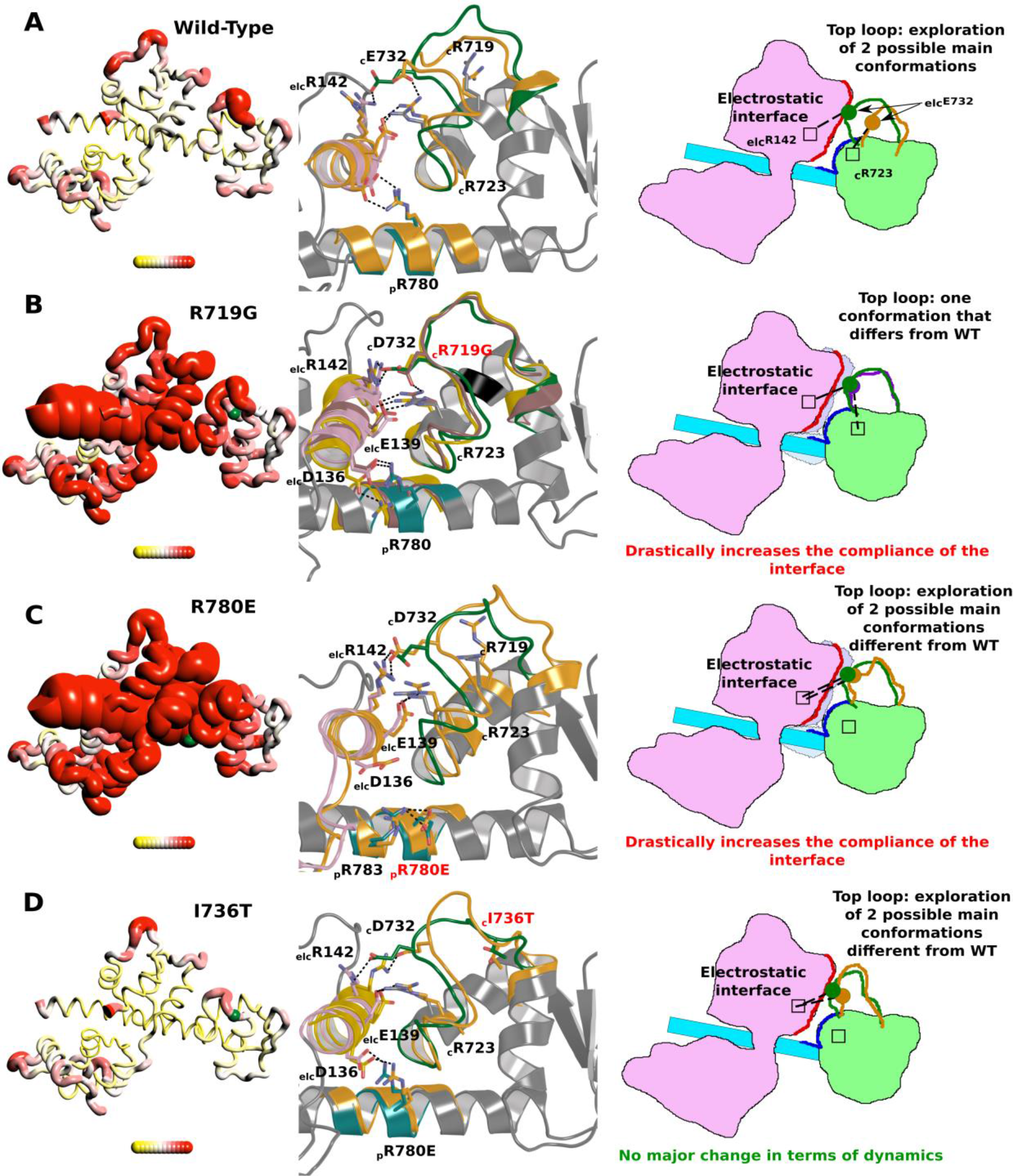
Dynamics of the Converter/ELC interface. Schematic representation of the results from the molecular dynamics simulations for the wild-type (WT), two proof-of-concept (R719A and R780E) mutations as well as one HCM (I736T) mutant that have been analyzed *in silico*. On the left, a “Putty representation” is displayed for a β-cardiac myosin construct that includes the residues 701-806 and the ELC. R.m.s. fluctuations during 30 ns simulations are represented with a r.m.s. scale ranging from 0.0621324 nm (in yellow) to 0.585496 nm (in red). Structures represented here correspond to the most populated structural state. In each structure, the position of the residue mutated are represented by a green sphere. On the center, a schematic representation of the region containing the converter, the ELC and the lever arm is displayed. The different populations of the top loop allowed by the dynamics of this region are drawn and the nature of the interactions between the converter and the ELC is also represented. On the right, the interface between the ELC, the converter and the pliant region is represented. The different positions schematized in the center are represented on the structure with the position of all key-residues that maintain the interface and its plasticity. **(A)** WT, **(B)** R719G, **(C)** R780E, **(D)** I736T.

**Supplementary Figure 3:**
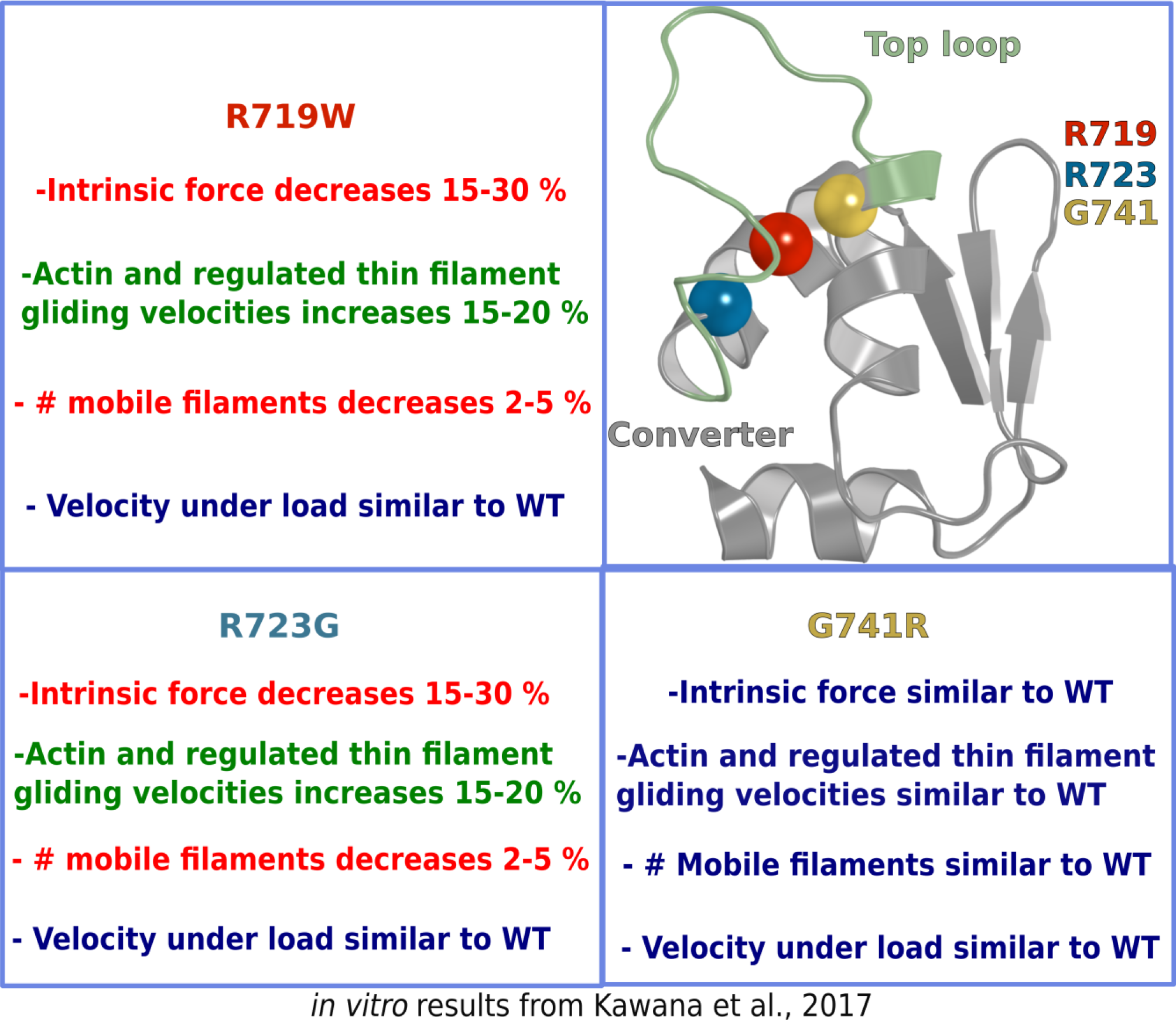
consequences of HCM mutations in the converter on the biomechanical properties of β-cardiac myosin as reported from the Kawana et al studies performed in the Spudich laboratory. Summary of the consequences of the mutations R719W, R723G and G741R compared to WT for the biomechanical properties of the cardiac myosin motor^31^. On the top, right, the three mutations are located (colored spheres) on a cartoon representation of the converter of β-cardiac myosin.

**Supplementary Figure 4:**
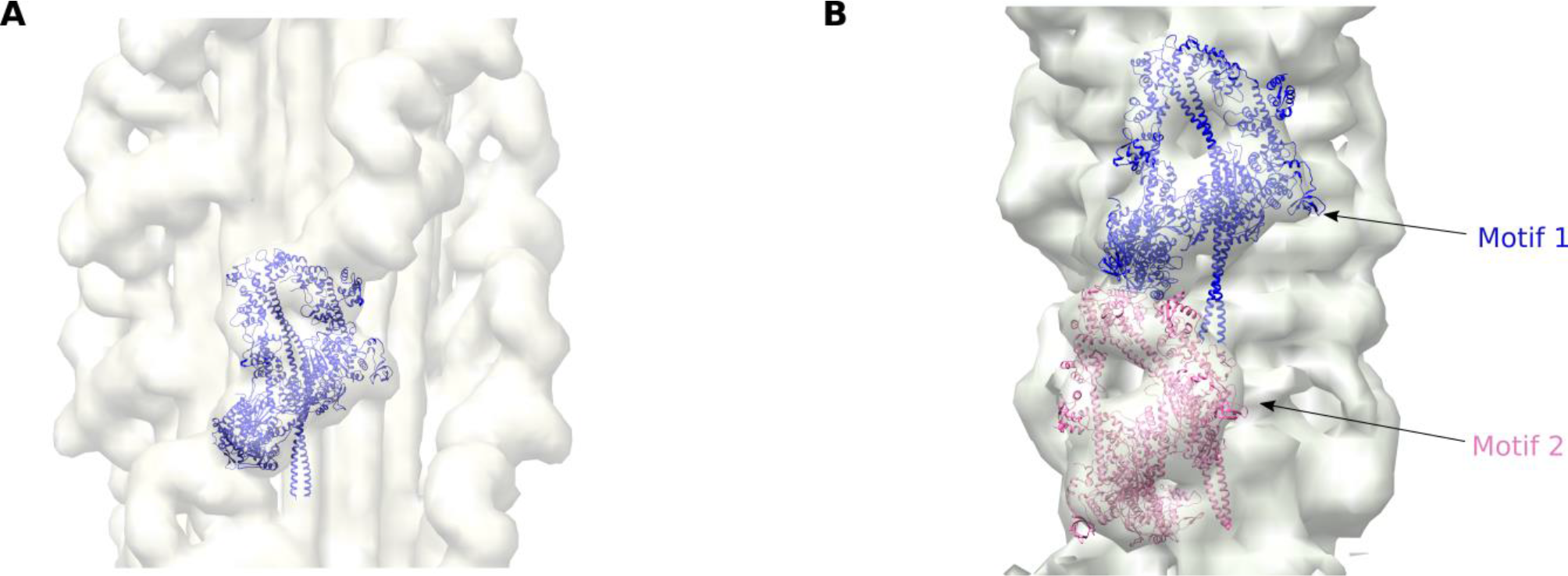
the sequestered state model and its fit in the electron density map. **(A)** Fit of our model in the electron density of tarantula muscle filament (EMD code: EMD-1950). **(B)** Reconstitution of the cardiac filament and sequestered state. Fit of our model in the human cardiac muscle filament electron density map obtained from negative stained EM data (EMDB code EMD-2240). Since the geometry in cardiac filament is not purely helical^23^, two motifs have been fitted in the filament, each motif is slightly rotated compared to the other.

**Supplementary Figure 5:**
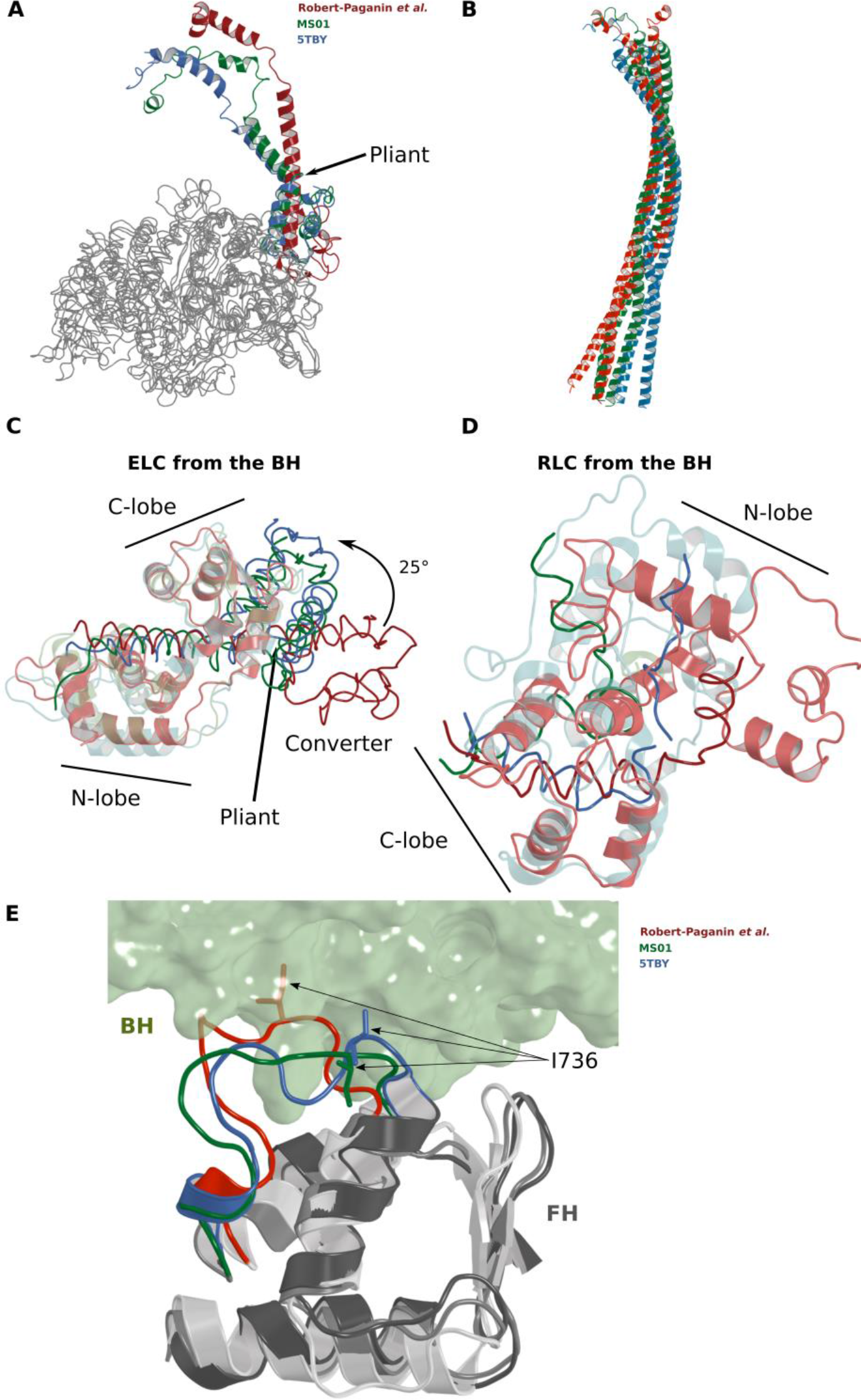
comparison of sequestered state models. Comparison of the sequestered model we propose (red) with the two previously released models: MS01 (green) from Jim Spudich’s lab^38^ and 5TBY (blue) from Raul Padron’s lab^30^. All the models have been superimposed on the N-ter region (1-171) of the blocked head (BH) of our model. **(A)** comparison of the converter/lever arm region that is displayed in cartoon models, **(B)** comparison of the S2 region. **(C)** Comparison of the converter/ELC region. **(D)** Comparison of the RLC region, RLC are aligned on the C-lobe. **(E)** Comparison of the interface between the FH-converter (cartoon) and the BH (transparent surface) in the model we present here (converter in white, top loop in red), the MS01 model (converter in light grey, top loop in green), and the 5TBY model (converter in dark grey, top loop in blue). For each structure, the side chain of Ile 736 is represented, showing that only in our model is this residue modeled in the interface between both heads.

**Supplementary Table 1:** Data collection and refinement statistics.

| **Bovine Cardiac Myosin S1, PR state** |  |
| --- | --- |
| Space group | P1 |
| Unit cell parameters | a = 55.118 Å<br>b = 94.319 Å<br>c = 125.411 Å<br>$\alpha = 104.76^\circ$<br>$\beta = 92.34^\circ$<br>$\gamma = 100.06^\circ$ |
| Source | ESRF (id 23-2) |
| wavelength (Å) | 0.87290 |
| Resolution (Å)<br>(outer shell) | 47.97 - 2.33<br>(2.413 - 2.33) |
| Reflections measured | 359796 (37143) |
| Unique reflections | 100862 (10032) |
| Redundancy | 3.6 |
| Completeness (%)<br>(outer shell) | 98.46 (97.87) |
| Rmeas (%) | 16.05 (135.20) |
| CC(1/2) | 99.4 (43.3) |
| $\langle I \rangle / \langle \sigma(I) \rangle$ | 7.76 (1.09) |
| **Refinement statistics** |  |
| Phasing | Molecular replacement |
| Molecules/UA | 2 |
| Rwork/Rfree (%) | 18.95/22.69 |
| Number of atoms non-H | 15508 |
| <B> (Å<sup>2</sup>) | 67.65 |
| r.m.s.d. bonds (Å) | 0.015 |
| r.m.s.d. angles (°) | 1.79 |
| **Ramachandran statistics** |  |
| Most favored (%) | 96.97 |
| Allowed (%) | 2.81 |
| Outliers (%) | 0.22 |

## References

1. Morita, H. et al. Shared Genetic Causes of Cardiac Hypertrophy in Children and Adults. N. Engl. J. Med. 358, 1899–1908 (2008).

2. Maron, B. J. & Maron, M. S. Hypertrophic cardiomyopathy. Lancet 381, 242–255 (2013).

3. Cirino, S., Lima, F. S. & Gonçalves, M. B. Spatial distribution of specialized cardiac care units in the state of Santa Catarina. Rev. Saude Publica 48, 916–924 (2014).

4. Kaski, J. P. et al. Prevalence of sarcomere protein gene mutations in preadolescent children with hypertrophic cardiomyopathy. Circ. Cardiovasc. Genet. 2, 436–441 (2009).

5. Garcia-Giustiniani, D. et al. Phenotype and prognostic correlations of the converter region mutations affecting the beta myosin heavy chain. Heart 101, 1047–1053 (2015).

6. Houdusse, A. & Sweeney, H. L. How Myosin Generates Force on Actin Filaments. Trends in Biochemical Sciences 41, 989–997 (2016).

7. Colegrave, M. & Peckham, M. Structural implications of beta-cardiac myosin heavy chain mutations in human disease. Anat Rec 297, 1670–1680 (2014).

8. Malik, F. I. et al. Cardiac myosin activation: A potential therapeutic approach for systolic heart failure. Science (80-. ). 331, 1439–1443 (2011).

9. Ochala, J. & Sun, Y. B. Novel myosin-based therapies for congenital cardiac and skeletal myopathies. J. Med. Genet. 53, 651–654 (2016).

10. Green, E. M. et al. Heart disease: A small-molecule inhibitor of sarcomere contractility suppresses hypertrophic cardiomyopathy in mice. Science (80-. ). 351, 617–621 (2016).

11. Planelles-Herrero, V. J., Hartman, J. J., Robert-Paganin, J., Malik, F. I. & Houdusse, A. Mechanistic and structural basis for activation of cardiac myosin force production by omecamtiv mecarbil. Nat. Commun. 8, 190 (2017).

12. Liu, C., Kawana, M., Song, D., Ruppel, K. M. & Spudich, J. A. Controlling load-dependent contractility of the heart at the single molecule level. (2018).

13. Köhler, J. et al. Mutation of the myosin converter domain alters cross-bridge elasticity. Proc. Natl. Acad. Sci. U. S. A. 99, 3557–3562 (2002).

14. Seebohm, B. et al. Cardiomyopathy mutations reveal variable region of myosin converter as major element of cross-bridge compliance. Biophys. J. 97, 806–824 (2009).

15. Brenner, B., Seebohm, B., Tripathi, S., Montag, J. & Kraft, T. Familial hypertrophic cardiomyopathy: Functional variance among individual cardiomyocytes as a trigger of fhc-phenotype development. Front. Physiol. 5, 392 (2014).

16. Houdusse, a, Szent-Gyorgyi, a G. & Cohen, C. Three conformational states of scallop myosin S1. Proc. Natl. Acad. Sci. U. S. A. 97, 11238–11243 (2000).

17. Spudich, J. A. The myosin mesa and a possible unifying hypothesis for the molecular basis of human hypertrophic cardiomyopathy. Biochem. Soc. Trans. 43, 64–72 (2015).

18. Homburger, J. R. et al. Multidimensional structure-function relationships in human beta-cardiac myosin from population-scale genetic variation. Proc Natl Acad Sci U S A 113, 6701–6706 (2016).

19. Trivedi, D. V., Adhikari, A. S., Sarkar, S. S., Ruppel, K. M. & Spudich, J. A. Hypertrophic cardiomyopathy and the myosin mesa: viewing an old disease in a new light. Biophys. Rev. (2017). doi:10.1007/s12551-017-0274-6

20. Wendt, T., Taylor, D., Trybus, K. M. & Taylor, K. Three-dimensional image reconstruction of dephosphorylated smooth muscle heavy meromyosin reveals asymmetry in the interaction between myosin heads and placement of subfragment 2. Proc. Natl. Acad. Sci. 98, 4361–4366 (2001).

21. Suk Jung, H. S., Komatsu, S., Ikebe, M. & Roger Craig, R. Head-Head and Head-Tail Interaction: A General Mechanism for Switching Off Myosin II Activity in Cells. Mol. Biol. Cell 19, 3234–3242 (2008).

22. Zoghbi, M. E., Woodhead, J. L., Moss, R. L. & Craig, R. Three-dimensional structure of vertebrate cardiac muscle myosin filaments. Proc Natl Acad Sci U S A 105, 2386–2390 (2008).

23. Al-Khayat, H. A., Kensler, R. W., Squire, J. M., Marston, S. B. & Morris, E. P. Atomic model of the human cardiac muscle myosin filament. Proc. Natl. Acad. Sci. U. S. A. 110, 318–23 (2013).

24. Alamo, L. et al. Conserved Intramolecular Interactions Maintain Myosin Interacting-Heads Motifs Explaining Tarantula Muscle Super-Relaxed State Structural Basis. J. Mol. Biol. 428, 1142–1164 (2016).

25. Hooijman, P., Stewart, M. A. & Cooke, R. A new state of cardiac myosin with very slow ATP turnover: A potential cardioprotective mechanism in the heart. Biophys. J. 100, 1969–1976 (2011).

26. Naber, N., Cooke, R. & Pate, E. Slow myosin ATP turnover in the super-relaxed state in tarantula muscle. J. Mol. Biol. 411, 943–950 (2011).

27. Stewart, M. A., Franks-Skiba, K., Chen, S. & Cooke, R. Myosin ATP turnover rate is a mechanism involved in thermogenesis in resting skeletal muscle fibers. Proc. Natl. Acad. Sci. 107, 430–435 (2010).

28. Wilson, C., Naber, N., Pate, E. & Cooke, R. The myosin inhibitor blebbistatin stabilizes the super-relaxed state in skeletal muscle. Biophys. J. 107, 1637–1646 (2014).

29. Irving, M. Regulation of Contraction by the Thick Filaments in Skeletal Muscle. Biophysical Journal 113, 2579–2594 (2017).

30. Alamo, L. et al. Effects of myosin variants on interacting-heads motif explain distinct hypertrophic and dilated cardiomyopathy phenotypes. Elife 6, 2386–2390 (2017).

31. Kawana, M., Sarkar, S. S., Sutton, S., Ruppel, K. M. & Spudich, J. A. Biophysical properties of human beta-cardiac myosin with converter mutations that cause hypertrophic cardiomyopathy. Sci Adv 3, e1601959 (2017).

32. Mura, C. Development & implementation of a PyMOL ‘putty’ representation. 3–5 (2004).

33. Nag, S. et al. The myosin mesa and the basis of hypercontractility caused by hypertrophic cardiomyopathy mutations. Nat. Struct. Mol. Biol. 24, 525–533 (2017).

34. Perrot, A. et al. Prevalence of cardiac beta-myosin heavy chain gene mutations in patients with hypertrophic cardiomyopathy. J. Mol. Med. (Berl). 83, 468–477 (2005).

35. Dominguez, C., Boelens, R. & Bonvin, A. HADDOCK: a protein-protein docking approach based on biochemical or biophysical Information. J. Am. Chem. Soc. 1731–1737 (2003). doi:10.1021/ja026939x

36. Llinas, P. et al. How Actin Initiates the Motor Activity of Myosin. Dev. Cell 33, 401–412 (2015).

37. Adhikari, A. S. et al. Early-Onset Hypertrophic Cardiomyopathy Mutations Significantly Increase the Velocity, Force, and Actin-Activated ATPase Activity of Human beta-Cardiac Myosin. Cell Rep 17, 2857–2864 (2016).

38. Nag, S. et al. Contractility parameters of human beta-cardiac myosin with the hypertrophic cardiomyopathy mutation R403Q show loss of motor function. Sci. Adv. 1, e1500511 (2015).

39. Mentes, A. et al. High-resolution cryo-EM structures of actin-bound myosin states reveal the mechanism of myosin force sensing. Proc. Natl. Acad. Sci. U. S. A. 115, 1292–1297 (2018).

40. Kampourakis, T., Zhang, X., Sun, Y.-B. & Irving, M. Omecamtiv mercabil and blebbistatin modulate cardiac contractility by perturbing the regulatory state of the myosin filament. J. Physiol. 596, 31–46 (2018).

41. Alamo, L., Pinto, A., Sulbarán, G., Mavárez, J. & Padrón, R. Lessons from a tarantula: new insights into myosin interacting-heads motif evolution and its implications on disease. Biophys. Rev. (2017). doi:10.1007/s12551-017-0292-4

42. Mijailovich, S. M. et al. Modeling the Actin.myosin ATPase Cross-Bridge Cycle for Skeletal and Cardiac Muscle Myosin Isoforms. Biophys. J. 112, 984–996 (2017).

43. Chaikuad, A., Knapp, S. & Von Delft, F. Defined PEG smears as an alternative approach to enhance the search for crystallization conditions and crystal-quality improvement in reduced screens. Acta Crystallogr. Sect. D Biol. Crystallogr. 71, 1627–1639 (2015).

44. Kabsch, W. XDS. Acta Crystallogr. D. Biol. Crystallogr. 66, 125–132 (2010).

45. McCoy, A. J. et al. Phaser crystallographic software. J. Appl. Crystallogr. 40, 658–674 (2007).

46. Emsley, P. & Cowtan, K. Coot: model-building tools for molecular graphics. Acta Crystallogr. D. Biol. Crystallogr. 60, 2126–2132 (2004).

47. Bricogne, G. et al. BUSTER version 2.10.2. Cambridge, United Kingdom Glob. Phasing Ltd. (2017).

48. Alamo, L. et al. Three-Dimensional Reconstruction of Tarantula Myosin Filaments Suggests How Phosphorylation May Regulate Myosin Activity. J. Mol. Biol. 384, 780–797 (2008).

49. Pettersen, E. F. et al. UCSF Chimera––a visualization system for exploratory research and analysis. J. Comput. Chem. 25, 1605–1612 (2004).

50. Jo, S., Kim, T., Iyer, V. G. & Im, W. CHARMM-GUI: a web-based graphical user interface for CHARMM. Journal of computational chemistry 29, 1859–1865 (2008).

51. Gilmore, J. H. NIH Public Access. North 29, 1883–1889 (2008).

52. van Zundert, G. C. P. et al. The HADDOCK2.2 Web Server: User-Friendly Integrative Modeling of Biomolecular Complexes. J. Mol. Biol. 428, 720–725 (2016).

53. Abraham, M. J. et al. Gromacs: High performance molecular simulations through multi-level parallelism from laptops to supercomputers. SoftwareX 1-2, 19–25 (2015).

54. Brooks, B. R. et al. CHARMM: the biomolecular simulation program. J. Comput. Chem. 30, 1545–1614 (2009).

